# Estimating the Readily-Releasable Vesicle Pool Size at Synaptic Connections in a Neocortical Microcircuit

**DOI:** 10.1101/646497

**Authors:** N Barros-Zulaica, J Rahmon, G Chindemi, R Perin, H Markram, S Ramaswamy, E Muller

**Affiliations:** Blue Brain Project, École polytechnique fédérale de Lausanne, Campus Biotech, 1202 Geneva, Switzerland; Blue Brain Project, École polytechnique fédérale de Lausanne, LNMC, 1015 Lausanne, Switzerland

**Keywords:** synaptic transmission, quantal analysis, multi vesicular release, neocortex, mathematical model, short-term depression

## Abstract

Previous studies based on the ‘Quantal Model’ for synaptic transmission suggested that neurotransmitter release is mediated by a single release site at individual synaptic contacts in the neocortex. However, recent studies seem to contradict this hypothesis and indicate that multi-vesicular release (MVR) could better explain the synaptic response variability observed *in vitro*. In this study we present a novel method to estimate the number of release sites per synapse, also known as the size of the readily-releasable pool (N_RRP_), from paired whole-cell recordings of layer 5 thick tufted pyramidal cell (L5_TTPC) connections in the somatosensory neocortex. Our approach extends the work of Loebel and colleagues to take advantage of a recently reported data-driven biophysical model of neocortical tissue. Using this approach, we estimated N_RRP_ to be between two to three for connections between L5-TTPC. To constrain N_RRP_ values for other connections in the microcircuit, we developed and validated a generalization approach using data on post-synaptic potential (PSP) coefficient of variations (CVs) from literature and matching to *in silico* experiments. Our study shows that synaptic connections in the neocortex generally are mediated by MVR and provides a data-driven approach to constrain the MVR model parameters of the microcircuit.

## 1 Introduction

Synaptic transmission is the basis for neuronal communication and information processing in the brain. Synaptic communication between neurons is mediated by neurotransmitters contained in presynaptic vesicles that are stochastically released from axonal boutons by incoming action potentials (APs) and diffuse across the synaptic cleft to bind receptors. Synaptic receptors are a class of ion channels which open as a result of transmitter binding, and the resulting transmembrane currents either depolarize or hyperpolarize the postsynaptic membrane, depending on the ion to which the channel is permeable (Mason et al., 1991; Südhof, 2000). Understanding the mechanisms behind vesicle release is crucial to unravel how information propagates between different types of neurons (Tsodyks and Markram, 1997). In fact, disruptions in vesicle release are implicated in pathologies such as Alzheimer’s disease or schizophrenia (Waites and Garner, 2011).

In 1954, del Castillo and Katz described the ‘Quantal model’ of synaptic transmission (Del Castillo and Katz, 1954). This model is characterized by the number of independent release sites (N), the probability of releasing a vesicle in the presynaptic cell followed by an AP (p) and the content of each vesicle, the quantal size (q), which collectively determine the efficacy of synaptic transmission (Del Castillo and Katz, 1954; Tsodyks and Markram, 1997). Previously, it was thought that no more than one vesicle could be released per synaptic contact, leading to the univesicular release hypothesis (UVR), in which N is equal to the number of physical synaptic contacts in a neuronal connection, at least for synapses in the neocortex (Biró et al., 2005; Korn et al., 1981, 1994; Silver et al., 2003). However, evidences as fluctuations of evoked post-synaptic potentials (PSPs) (Tang et al., 1994), large concentration of neurotransmitter in the synaptic cleft (Tong and Jahr, 1994) or a high range variability of receptor-mediated signals of N-methyl-D-aspartate (NMDA) and α-amino-3-hydroxy-5-methyl-4-isoxazolepropionic acid (AMPA) receptors (Conti and Lisman, 2003) suggested that transmission at a single synaptic contact could be multiquantal. Consequently, a multivesicular release hypothesis (MVR) was proposed, where several release sites could underlie a synaptic contact in a neuronal connection. In fact, there are evidences showing that MVR occurs in brain regions such as the hippocampus (Christie and Jahr, 2006; Tong and Jahr, 1994), the cerebellum (Auger et al., 1998), the hypothalamus (Gordon, 2005) or the cerebral cortex (Huang et al., 2010; Molnár et al., 2016; Rudolph et al., 2015). Furthermore, MVR has been physically observed in individual contact sites of the mouse hippocampus with ‘flash-and-freeze’ electron microscopy technique (Watanabe et al., 2013).

In the neocortex, recent studies in rodents support the idea of MVR between pyramidal cells (Hardingham et al., 2010; Loebel et al., 2009; Rollenhagen et al., 2018). It has also been reported that different cortical areas show different vesicular release behavior. For instance, it has been reported that connections between cells in layer 4 exhibit UVR in the primary visual cortex, but the primary somatosensory cortex exhibit MVR (Huang et al., 2010). By contrast, other studies have reported that connections between layer 4 stellate cells and layer 2/3 pyramidal cells in the rat barrel cortex (Silver et al., 2003) and between pyramidal cells and interneurons in the rat cortex (Molnár et al., 2016) follow the UVR hypothesis. Moreover, in human neocortex, MVR has been observed for connections between pyramidal cells and fast spiking interneurons (Molnár et al., 2016).

Theoretical models have also contributed to understand vesicular release by studying synaptic processes as short-term synaptic plasticity. These models account for crucial parameters for modeling the presynaptic release such as the number of vesicles available for release and the probability of releasing these vesicles (Hennig, 2013; Loebel et al., 2009; Tsodyks and Markram, 1997). These models also assume that the rate of releasable neurotransmitter is limited and they take into account vesicle depletion (Liley and North, 1953), facilitation mechanisms (Betz, 1970; Markram et al., 1998; Varela et al., 1997) and vesicle replenishment (Fuhrmann et al., 2004). Many models has reported evidences of the importance in the number of release sites per neuronal connection for instance for information coding (Fuhrmann et al., 2002). It has also been reported that the frequency in which multiple vesicles are released and the number of vesicles released is very important for receptor activation (Boucher et al., 2010). Some studies also reveal the importance of having the number of readily releasable pool (N_RRP_) higher than 1 for being able to reproduce plasticity processes (Nadkarni et al., 2010). N_RRP_ represents the number of docked vesicles ready to be released at boutons of the presynaptic axon when an action potential is propagating. Other works found evidences of MVR at different synapses (Loebel et al., 2009) using synaptic models.

As MVR is a complex process that is suspected to regulate synaptic transmission and plasticity by increasing the dynamic range of synapses (Fuhrmann et al., 2004) influencing cognitive functions such as learning and memory. MVR is thought to directly impact synaptic noise and variability, with larger vesicle pool sizes resulting in a reduction of synaptic noise, with important implications for the transmission of information (Fuhrmann et al., 2004). Thus, accurate models of synaptic transmission in the neocortical microcircuit (Markram et al., 2015) must therefore account for the current state of knowledge on MVR.

Taken together, these previous studies demonstrate the complexity of the vesicle release process in the cortex, and show that there remains a need for developing an integrated consensus on vesicle pool sizes in the rodent neocortex.

In this work, we estimated the average size of the N_RRP_ for individual synaptic contacts between different cell types in the neocortical microcircuit. To compute this parameter, we employed data-driven simulations of connected pairs of neurons (Markram et al., 2015) with previous reported counterpart *in vitro* experiments, taking care to reproduce pertinent aspects of the experiments such as the spatial distributions of neuron sampling, specific synaptic parameters and sub-threshold noise sources. We then optimized model MVR parameters, in particular N_RRP_, to reproduce response variability as observed in the experiments, as assessed by coefficients of variation (CVs) of excitatory postsynaptic potentials (EPSPs). We first made use of a high quality *in vitro* dataset on synaptic connections between layer 5 thick-tufted pyramidal cells (L5_TTPC) in the juvenile rat somatosensory cortex to estimate synaptic and noise model validity and the MVR free parameter N_RRP_, extending the work of Loebel and colleges (Loebel et al., 2009) to morphologically detailed neuron models with anatomically and physiologically constrained connectomes. We further developed an approach to estimate N_RRP_ for both excitatory and inhibitory connection types using literature data sources providing PSP CVs. Our work provides evidence for MVR at most of the connection types examined, albeit with lower N_RRP_ values than previously reported by Loebel et al., 2009, suggesting it could be a general property of local cortical connections.

## 2 Materials and Methods

### 2.1 Slice preparation and electrophysiology

Fourteen- to eighteen-day-old Wistar rats were quickly decapitated according to the Swiss Welfare Act and the Swiss National Institutional Guidelines on Animal Experimentation for the ethical use of animals. The project was approved by the Swiss Cantonal Veterinary office following its ethical review by the State Committee for Animal Experimentation. Multiple (6-12 cells simultaneously) somatic whole cell patch-clamp recordings were carried out with Multiclamp 700B amplifiers in current clamp mode. Brain sagittal slices of 300 μM width were cut on an HR2 vibratome (Sigmann Elektronik). Temperature was held at 34 ± 1 °C in all experiments. The extracellular solution contained 125 mM NaCl, 2.5 mM KCl, 25 mM D-glucose, 25 mM NaHCO3, 1.25 mM NaH2PO4, 2 mM CaCl2, and 1 mM MgCl2 bubbled with 95% O2 and 5% CO2. The intracellular pipette solution contained 110 mM potassium gluconate, 10 mM KCl, 4 mM ATP-Mg, 10 mM phosphocreatine, 0.3 mM GTP, 10 Hepes, and 13 mM biocytin adjusted to pH 7.3–7.4 with 5 M KOH.

Data was acquired using PulseQ, a custom-writen data acquisition library written in IGOR (WaveMetrics, Lake Oswego, OR). L5_TTPCs were selected according to their large soma size (15–25 μm) and their apparent large trunk of the apical dendrite. Cells were visualized by infrared differential interference contrast video microscopy using a VX55 camera (Till Photonics) mounted on an upright BX51WI microscope (Olympus). Data acquisition was performed through an ITC-1600 board (Instrutech) connected to a PC running a custom-written routine (PulseQ) under IGOR Pro (Wavemetrics). Sampling rates were 5-10 kHz, and the voltage signal was filtered with a 2-kHz Bessel filter. The stimulation protocol consisted of pre-synaptic stimulation with eight electric pulses at 20 Hz followed by a single pulse 500 ms later (recovery test), at the sufficient current intensity to generate APs in the presynaptic neuron while the postsynaptic neuron responses were recorded. The protocol was repeated from 20 to 60 times. (Fig.1A up)

**Figure 1.**
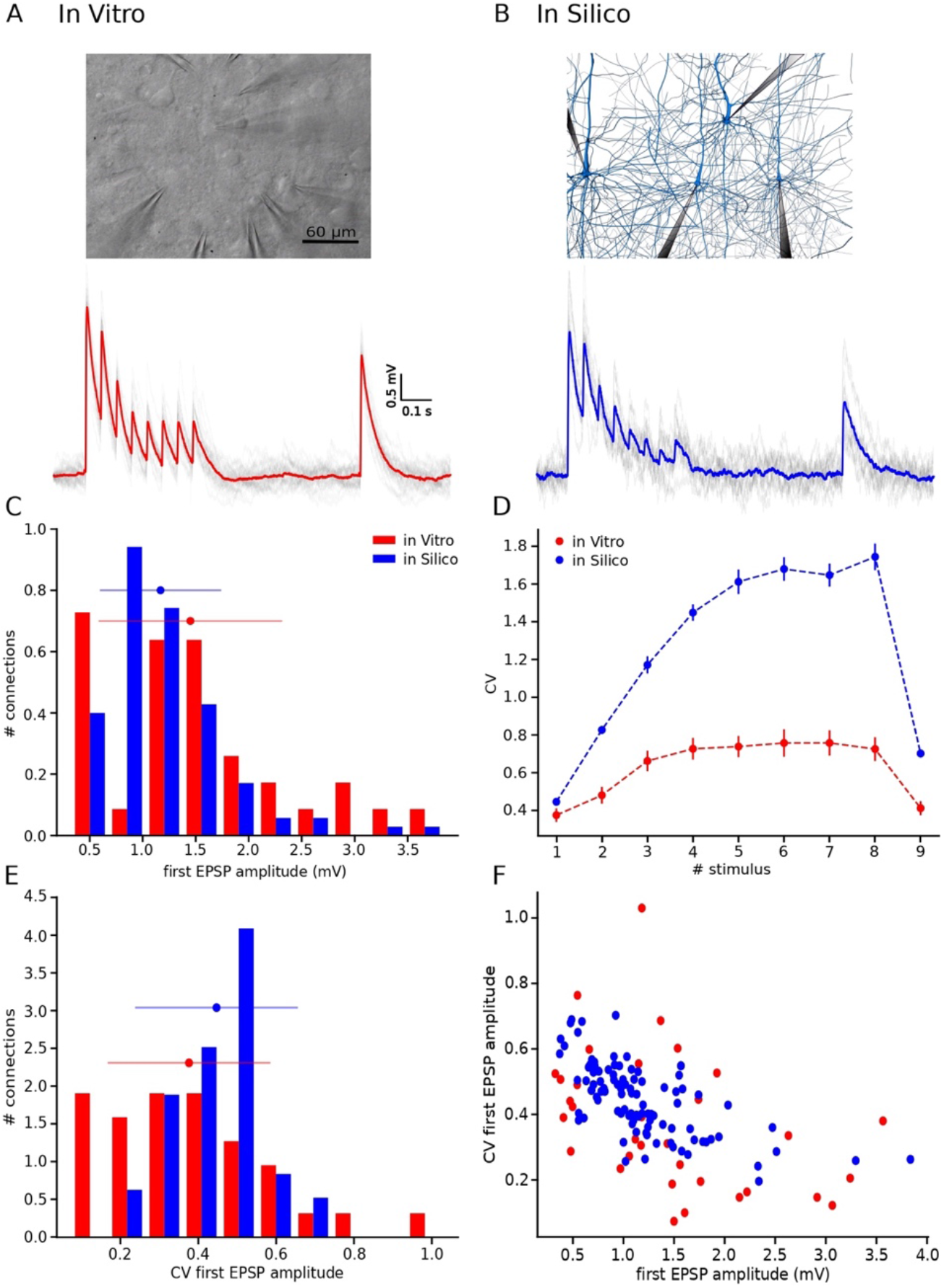
With the UVR hypothesis it was not possible to reproduce the variability observed *in vitro*. (**A**)(up) Picture of a multiple whole cell patch-clamp recording in L5_TTPC connections. (down) *In vitro* mean voltage trace (red) of 20 protocol repetitions (grey). **(B)**(up) Picture of a patch-clamp *in silico* experiment performed on L5_TTPC connections from the data-driven model of the rat cortex microcolumn. (down) *In silico* mean voltage trace (blue) of 20 protocol repetitions (grey). **(C)** Histogram showing the distribution of the first EPSP amplitude for *in vitro* (red) and for *in silico* (blue) experiments. **(D)** Mean CV profiles for the *in vitro* (red) and the *in silico* (blue) experiments. **(E)** CV distribution of the first EPSP amplitude for *in vitro* (red) and *in silico* (blue) data sets. **(F)** Raster plot of the first EPSP amplitude against the CV of the first EPSP amplitude for *in vitro* (red) and *in silico* (blue) experiments. In the distributions and the CV profile, dots represent the mean and vertical and horizontal bars represent the standard deviation of all the experiments respectively.

### 2.2 Stochastic model for short-term dynamics and multi-vesicular release

Our model describes the short-term synaptic dynamics defined by a stochastic generalization of the Tsodyks-Markram model (TM-model) (Maass and Markram, 2002; Tsodyks and Markram, 1997) that is known to fit excitatory as well as inhibitory synapses behavior of biological experiments (Gupta et al., 2000; Markram et al., 1998). This model considers that there is a finite number of vesicles ready to be released defined by N_RRP_ that could be in ready or recovery state. In this study we followed the synaptic dynamics described on Maass and Markram, 2002 that is able to predict the sequence of postsynaptic potential (PSP) amplitudes produced by any spike train. This behavior is described by four main synaptic parameters: the absolute synaptic efficacy (A), the fraction of synaptic resources used by a single spike (U), the time constant for recovery from facilitation (F) and the time constant for recovery from depression (D). The PSP amplitudes prediction obeys the following mathematical expressions:

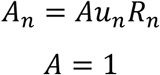

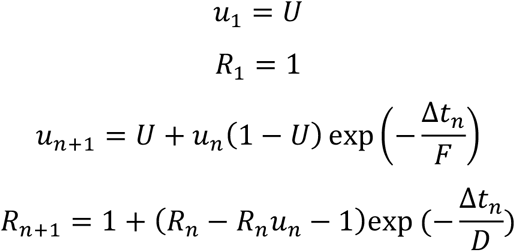

In short, when the nth spike occurs there is certain fraction of synaptic efficacy modeled by R_n_. Accordingly, the product u_n_R_n_ models the fraction of synaptic efficacy used by the nth spike. Combining these terms is possible to describe the fraction of synaptic efficacy available when the next spike arrives at time Δt_n_ assuming that the synaptic efficacy has an exponential recovery with time constant D. How much fraction of synaptic efficacy (R_n+1_) is used when (n+1)th spike occurs is defined by u_n+1_ which increases for each subsequent spike from u_n_ to U(1 – u_n_) + u_n_ and goes back to U following an exponential with time constant F (Maass and Markram, 2002).

Our model also takes into consideration postsynaptic models for AMPA and NMDA receptors. Thus, if a vesicle is successfully released, these receptors get activated with a conductance g_max_/N_RRP_ with g_max_ as the maximal conductance.

### 2.3 Fitting synapse model parameters to the data

We constrained our model by extracting the parameters U, D and F from each *in vitro* connection (Fig.2C, D and E). For this purpose, we needed to compute the EPSP amplitudes of each averaged voltage trace. All experimental traces were normalized to their maximum value, so the maximum amplitude would be 1 and we could directly compute the peak value instead of the total amplitude. To perform an accurate computation of the peaks we used a mathematical tool for deconvolving the voltage averaged trace (Richardson and Silberberg, 2008). Basically, what this method provided us was the possibility of removing the smoothing effect of the membrane cell low-pass filter with a time scale equal to τ_mem_, so we could extract the peaks from the EPSPs, all of them leveled to the same voltage base (Fig.2B).

**Figure 2.**
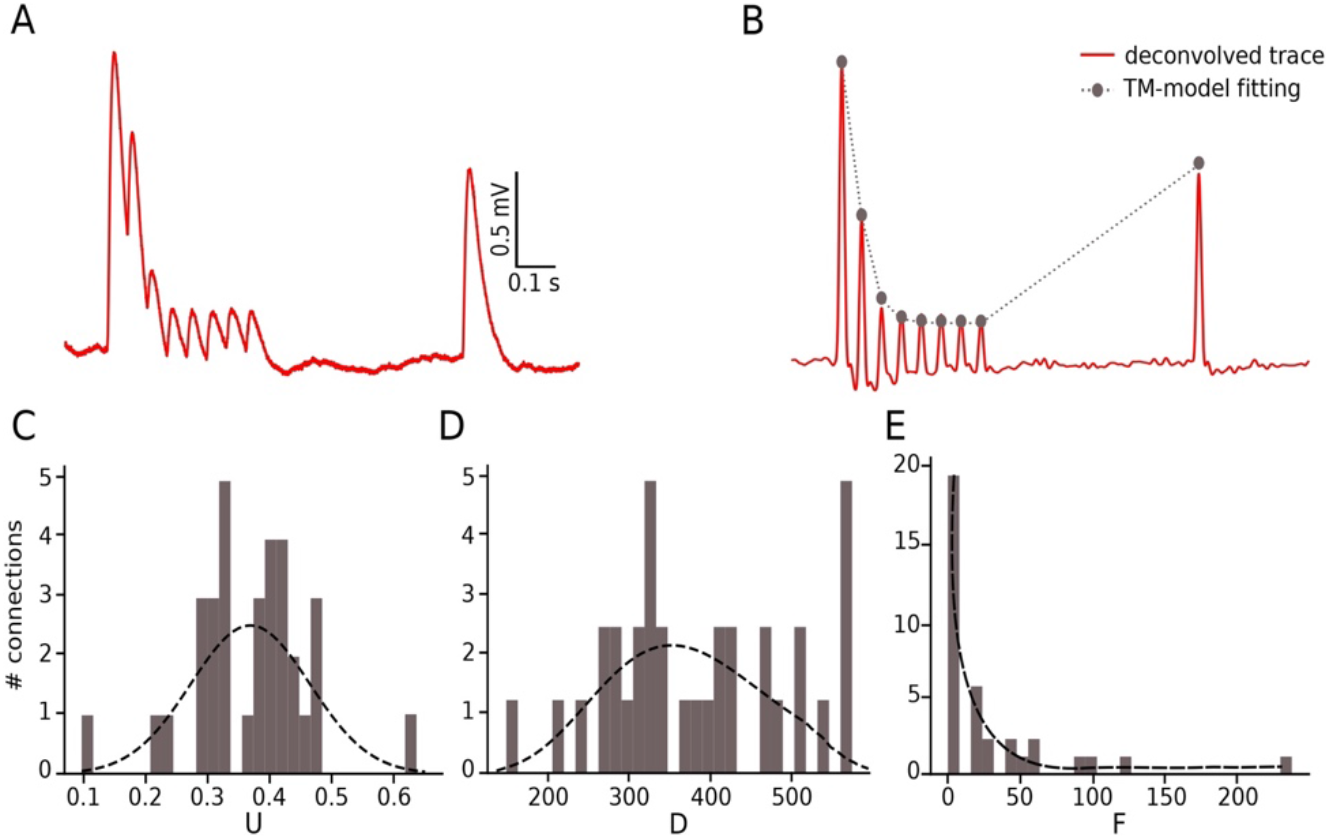
Fitting in vitro data to the TM-model. **(A)** Example of an *in vitro* mean voltage trace of L5_TTPC connection. **(B)** Corresponding deconvolved voltage trace (red) with the fit to the deterministic TM-model (grey). **(C)** Distribution of the probability of release parameter (U), **(D)** distribution of the time to recovery from depression (D) and **(E)** distribution of the time to recovery from facilitation (F). Values obtained from the fitting to the TM-model of 33 *in vitro* connections.

To express this process mathematically we used the next equation:

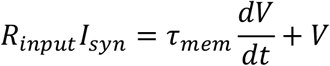

The right-hand part of the expression is the voltage deconvolution, while the left hand contains the unfiltered synaptic current. The requirement here is to compute τ_mem_ for each *in vitro* connection by fitting the decay part of the recovery peak (9^th^ EPSP) of the averaged voltage trace to an exponential.

Once we got the peaks, we matched the possible U, D and F parameters with the help of a genetic algorithm (Goldberg and Holland, 1988) with 500 generations. Using the mathematical expression of the model, the input values were the peaks of each trace and the ranges for U, D and F were (0, 1.0), (0, 1000.0) and (0, 2000.0) respectively. We obtained one set of U, D and F per connection and consequently we defined these parameters as distributions (normal, gamma and gamma respectively) with their correspondent mean and standard deviation.

### 2.4 In Silico experiments: the cortical microcircuit

For the *in silico* experiments we took advantage of the somatosensory cortical microcircuit built into the frame of the Blue Brain Project (Markram et al., 2015). This is a data driven model with detailed anatomy and physiology with 31000 neurons, 8 million connections and 37 million synapses in a volume of 0.29 mm^3^.

Once we got the values for the mean and the standard deviation of the synaptic parameters from the fitting of the *in vitro* data to the TM-model, we updated the simulation with these new values implemented as distribution defined by their mean and standard deviation. We also computed g_max_ through the simulation of synaptic connections with different g_max_ values and we kept the g_max_ values which first EPSP amplitude was similar to the real data. After all the updates, we were able to perform patch-clamp *in silico* experiments (Fig1.B up), with the same conditions as the *in vitro*, with different N_RRP_ values. These values were shaped defining the mean of a Poisson distribution of 100 points shifted one unit to the right. The range of means of the poissonian distributions varied from 0 to 13 (1 ≤ N_RRP_ ≥ 14) in the case of studying MVR and 0 (N_RRP_ = 1) while studying UVR. We decided to set the maximum value to 14 vesicles on average per release site because is already the double of what Loebel and colleges predicted on their research (Loebel et al., 2009).

Subsequently, we run simulations for 100 L5_TTPC connections of the microcolumn with 20 protocol repetitions. As these simulations will be used to be compare with the *in vitro* traces, we had to be sure that U, D and F distributions for the *in silico* connections had the same shape as for the *in vitro*. To this end we set the distributions before running the simulations. After running the simulations, we checked that the 1^st^ EPSP amplitude for the simulations were within the range of the *in vitro* data collection. In the case of UVR study we didn’t have to remove any experiment, but in the case of MVR we removed 15 connections (85 out of 100) according to the criteria explained before.

### 2.5 Noise calibration. Ornstein-Uhlenbeck process

After performing the simulations with different N_RRP_ values and selecting the simulations within the 1^st^ EPSP amplitude range of the *in vitro* connections, we had to complete the simulated traces with voltage fluctuations that will account for the membrane noise. One way of doing it was implementing the Ornstein-Uhlenbeck process (OU-process), which is a stochastic process that allowed us to simulate small random variability. The OU-process describes the velocity of the movement of a Brownian particle considering the friction and is a stationary Gauss-Markov process (Enrico Bibbona, 2008).

Mathematically the expression used in this work for this process was:

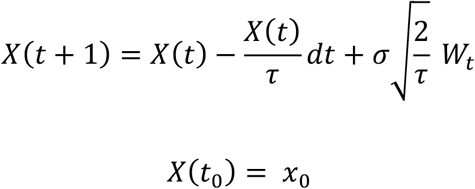

Where τ is the membrane time constant, σ is the standard deviation of the voltage and W_t_ is a random term coming from the Wiener process. In the case of σ = 0 the equation will have the solution *X*(*t*) = *x*_0_*e*^−(*t* − *t*_0_)/τ^ so X(t) relaxes exponentially towards 0. In general, X(t) fluctuates randomly, the third term pushes it away from zero, while the second term pulls it back to zero (Bradley Efron and Robert J. Tibshirani, 1994). In Physics this process is used to describe noisy relaxation activity.

In our specific case we defined σ and τ using the voltage values between the 8th and the 9th EPSPs, 400 ms in total, for each repetition (sweep) in a connection and then we averaged the resulting values (Fig.3A). By computing the standard deviation of these points, we got one σ per connection (33 in total). Through the calculation of the autocorrelation of this section and fitting it to an exponential, we got one τ per connection (Fig.3B). The computation of their means gave us the values to implement the membrane noise for the *in silico* traces (Fig.3C).

**Figure 3.**
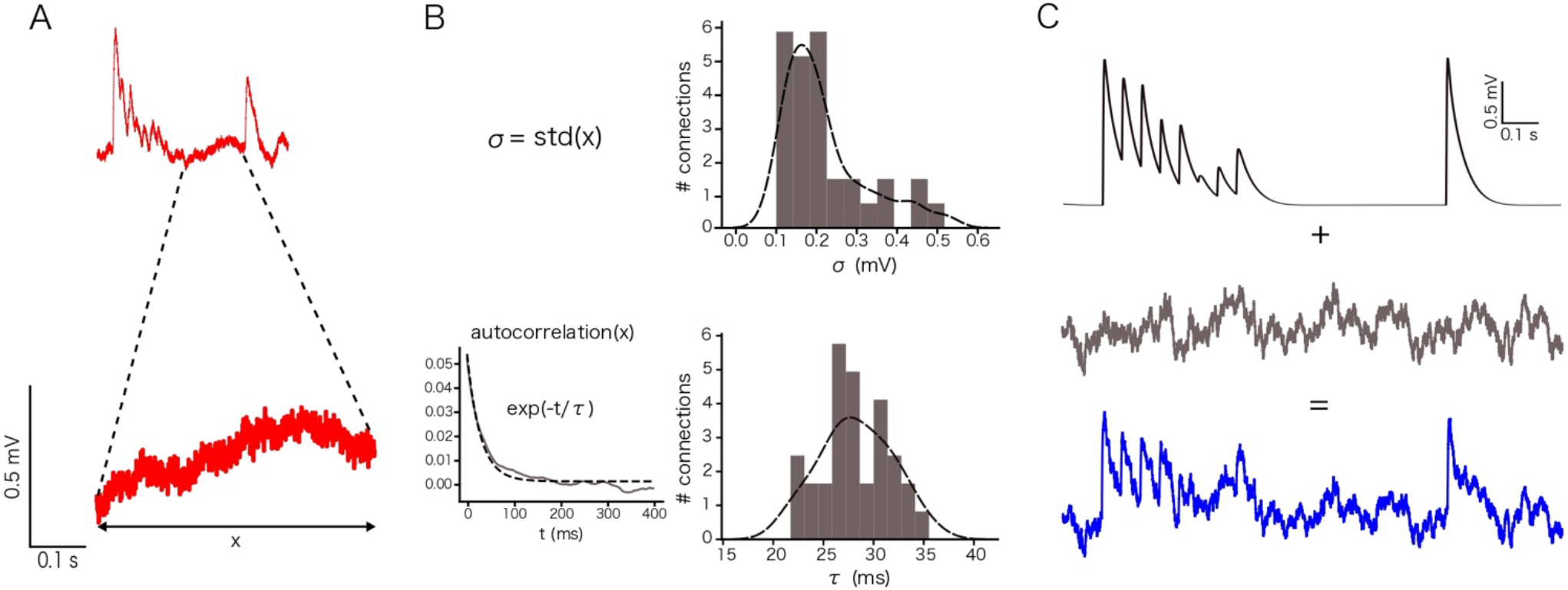
Noise calibration. **(A)** Example of an *in vitro* single protocol repetition (up). (down) Zoom over 400 ms segment used to compute the parameters for noise calibration. **(B)** Distribution of σ (up) and τ (down). σ was computed as the standard deviation of the voltage segment. τ was computed by fitting the voltage segment autocorrelation to an exponential. The distributions show the mean value s for the 33 *in vitro* connections. **(C)** (up) Single *in silico* trace without noise, (middle) OU-process generated to be added to the single *in silico* trace and (down) the noisy single protocol repetition that is the result of adding the previous two traces.

### 2.6 CV profile computation. The Jack-Knife bootstrapping analysis

In order to compute the CV for the EPSP amplitudes in the case of the *in vitro* and the *in silico* experiments in the more accurate possible way, we implemented the Jack-Knife method (JKK) (Bradley Efron and Robert J. Tibshirani, 1994).

This method consists in excluding one observation at a time from a group of observations. In our specific case, from a set of single traces we computed the average of all but one off the traces each time, obtaining at the end a set of averaged-JKK traces. From each of this averaged-JKK traces we computed the amplitudes for the nine EPSPs. This computation is much more precise considering we removed the noise by averaging. Then we could compute the CV profiles for the *in vitro* and the *in silico* experiments using the following equations:

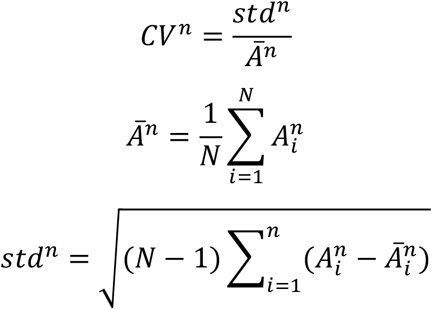

Where n denotes the EPSP index (n = 1,…,9) and N is the number of single traces per connection.

Once having both simulation sets, to study UVR and MVR, we could compute the CV profile for the EPSP amplitudes using the JKK approach in both cases and compare them with the CV profile computed with the *in vitro* data. We computed the mean square distance in order to obtain the minimum error between *in vitro* and *in silico* CV profiles (Fig. 4E). We iterated this procedure 50 times and then we provided the mean and the standard deviation for the N_RRP_ that correspond with the smallest error.

**Figure 4.**
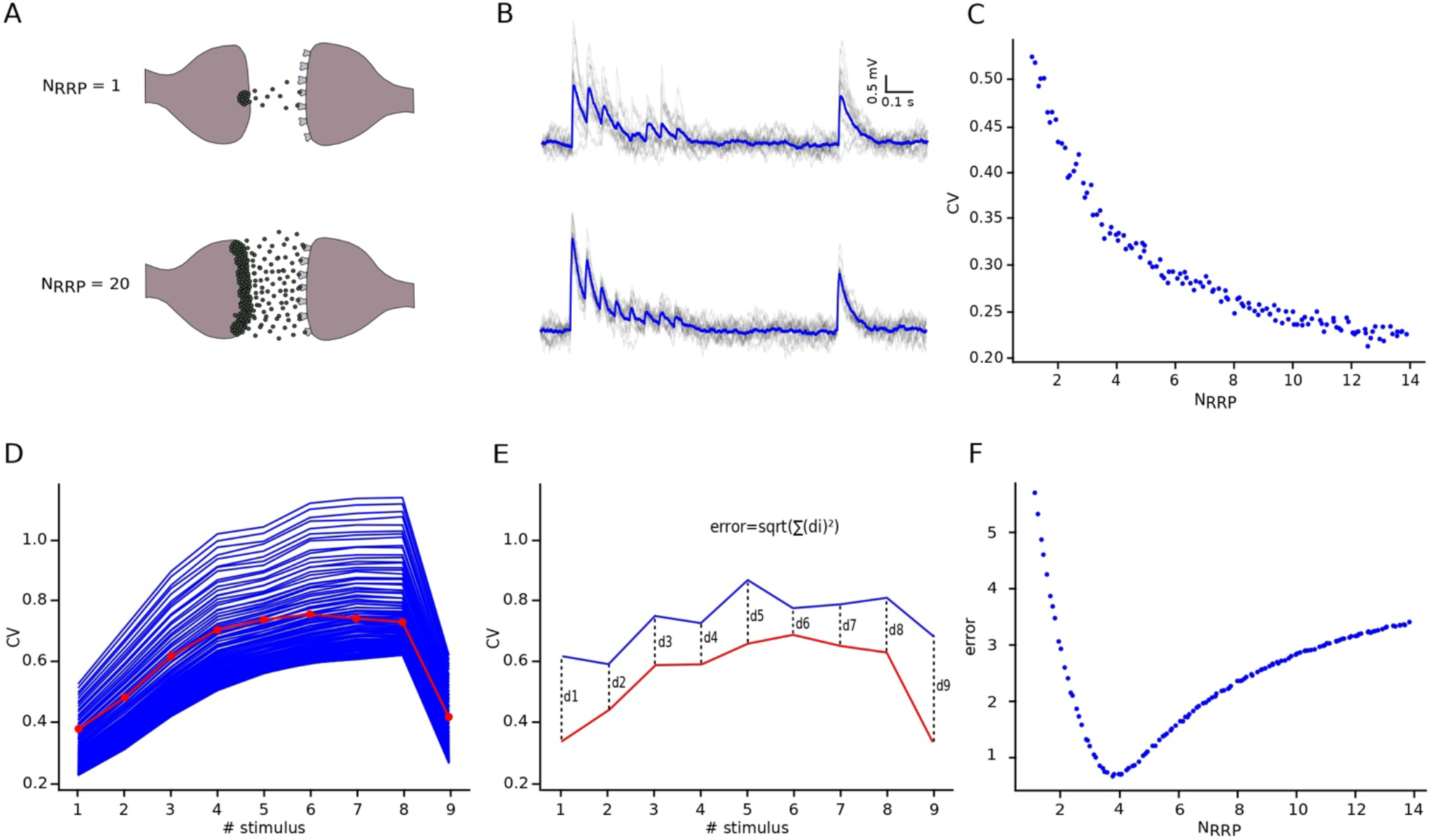
NRRP computation. **(A)** Illustration showing one synaptic connection releasing neuro transmitters from only one vesicle (up) and the same synaptic connection releasing neurotransmitters from twenty vesicles (down). **(B)** The corresponding effect of releasing neurotransmitters from one (up) or from twenty (down) vesicles reflected on the variability and shape of the *in silico* traces. The mean voltage traces are painted in blue while each protocol repetition is represented in grey. **(C)** Diagram showing the effect of N_RRP_ over the CV. **(D)** Mean CV profile for the *in vitro* (red) and all the *in silico* connections with different N_RRP_ values. **(E)** Diagram explaining the mean square distance computation. **(F)** N_RRP_ against error, showed a clear minimum around the value obtained for this specific connection.

## 3 Results

### 3.1 Motivation for implementing MVR in the model

Our simulation was based on a biologically detailed, data-driven model of a rat neocortical microcircuit (Markram et al., 2015). Although this data-driven simulation was capable of reproducing many biological results, some improvements were needed. For instance, our goal in this work, which was to be able to reproduce the synaptic release variability observed *in vitro*.

We began by implementing UVR at all synaptic contacts in the neocortical microcircuit model. As a result, our synaptic responses were highly variable in comparison against biological experiments. In order to study this problem, we recorded new 33 pairs of L5_TTPC strongly connected cells (Fig.1A up) and computed the amplitude and the CV of the amplitudes for each EPSP. In figure 1, we show two examples of experiments, one *in vitro* (Fig. 1A down red) and another *in silico* (Fig. 1B down blue). Differences in the shape and amplitude of the mean traces can be seen. The *in vitro* trace in red has a higher amplitude than the *in silico* in blue. Is also visible that the shape of the *in silico* mean trace is noisier than the *in vitro*, reflecting larger variability between protocol repetitions.

Although is possible to see differences between the amplitudes in the examples, the Kruskal-Wallis test of the distributions for the first EPSP amplitude (Fig. 1C) revealed no significant difference between the two data sets (mean ± standard deviation EPSP values: 1.46 ±0.86 mV for *in vitro*; 1.17 ± 0.57 mV for *in silico*; p = 0.15) (significant if p < 0.05). However, the computation of the Kruskal-Wallis test between the distributions of the CV for the first EPSP amplitude (Fig. 1E) revealed a significant difference between both data sets (mean CV values: 0.38 ± 0.21 for *in vitro*; 0.45 ± 0.11 for *in silico*; p = 0.0092). Consequently, computing the CV profile for the EPSP amplitudes for every stimulus showed a large difference between both data sets (Fig. 1D). The previous statistic test computed for the CV of the EPSP amplitudes showed very significant differences (p < 10^−9^) for the rest of the EPSPs. The distributions (Fig. 1C and E) were normalized to the respecting sample size such that the sum of products of width and height of each column is equal to the total count (33 for *in vitro*, 100 for *in silico*). Moreover, the cross validated Kolmogorov-Smirnov test for two-dimensional data (Press and Teukolsky, 1988) showed a significant difference between the first EPSP amplitude against the CV of the first EPSP amplitude for each data set (p = 0.0022; significant if p < 0.2) (Fig. 1F) demonstrating that both data sets are different. As we can observe in figure 1F, *in vitro* data (red dots) are more dispersed than *in silico* (blue dots).

This evidence motivated us to implement the MVR hypothesis as it is known to provide the synapse with a larger range of variability, from too variable to almost no variance (Brémaud et al., 2007; Wang et al., 2006).

### 3.2 Extracting values for the TM-model and noise calibration

To be able to compare the *in vitro* and the *in silico* experiments in a very precise manner, we had to extract certain parameters from the 33 *in vitro* traces mentioned before. The parameters in which we were interested were those related with the TM-model, to describe the synapse behavior, and the ones related with voltage fluctuations as the membrane noise.

The important synaptic parameters for the TM-model were extracted from the deconvolution of *in vitro* each averaged trace (Fig. 2B), so we were able to extract the values of the peaks from the same voltage level. We obtained three distributions, one per each parameter. For U we obtained a normal distribution with a mean value of 0.38 ± 0.1 (Fig. 2C), D fitted a gamma distribution with a mean value of 365.6 ± 100.15 ms (Fig. 2D) and F was also fitted to a gamma distribution with mean 25.71 ± 45.87 ms (Fig. 2E). These values were similar to the values found in previous studies (Tsodyks and Markram, 1997b; Wang et al., 2002). Later we implemented these parameters as the correspondent specific distributions into our model. The next step was to estimate the g_max_. To approximate it, we ran simulations with different g_max_, computed the amplitude of the first EPSP and compared this value with the values from the *in vitro* data set. We kept the g_max_ from which the correspondent EPSP amplitude was similar to the one found in the *in vitro*. The value we obtained was 1.54 ± 1.20 nS, which is in accordance with values found in previous works (Markram et al., 1997). These parameters helped us to simulate the synaptic behavior of L5_TTPC connections. (Values expressed by mean ± standard deviation).

Other important parameters to be calibrated are the ones related with membrane noise. For this purpose, we implemented an OU-process, for which we needed to extract some parameters from each individual sweep of the *in vitro* traces. The values that we got from this calibration were σ = 0.22 ± 0.10 (Fig. 3B up) and τ = 28.2 ± 3.5 (Fig. 3B down). After that we added synthetic voltage noise to each of the simulated sweeps generated each time as a new OU-process (Fig. 3C). With all these: the synaptic U, D, F and g_max_ and the membrane noise parameters, we captured the synaptic variability for this specific connection.

### 3.3 Optimizing N_RRP_ for L5_TTPC connections

After defining all the needed parameters, we could run simulations with different N_RRPs_ with values between 1 to 14 (see methods 2.4) and compare them against the CV obtained from the *in vitro* data computed through the JKK approach. We could observe the relation between N_RRP_ and the CV for this specific connection (Fig. 4C). As a first result we could observe that the CV for the first EPSP amplitude was higher when N_RRP_ was smaller. Therefore, for UVR like connections the variability between sweeps is larger than for MVR like connections. This result was in agreement with previous studies (Brémaud et al., 2007; Wang et al., 2006) and is also reflected on the *in silico* experiments run on the microcircuit with N_RRP_ = 1 (Fig. 4B, up) and N_RRP_ = 20 (Fig. 4B, down) used as examples to show how the variability and shape of the connections changes with the number of released vesicles.

In order to determine N_RRP_, we computed the CV profiles of the *in silico* experiments performed with different N_RRPs_ then by measuring the mean square distance (Fig. 4E) for each of them against the *in vitro* CV profile (Fig. 4D), we found that for this specific cell connection the minimum error was achieved by the simulation with N_RRP_ = 3.78 ± 1.65. This result showed us that MVR is a process that happens in the rat cortex between L5_TTPCs and is also in agreement with other studies (Loebel et al., 2009a; Rollenhagen et al., 2018).

### 3.4 Implementing MVR improved the variability of the synapses in the model

Once we obtained the value for the N_RRP_ parameter we wanted to compare the previous result in which only one vesicle was released (Fig. 1) and observe if the implementation of the MVR hypothesis really improved the model behavior, showing a result closer to the biological data. For this reason, we computed the distributions for the first EPSP amplitude, the CV of the first EPSP amplitude and the CV profile of the EPSP amplitudes for all the pulses. We could observe that the shape and the amplitude of the *in silico* trace (Fig. 5B) is similar to the *in vitro* trace (Fig. 5A), in contrast to the comparison with the UVR like *in silico* trace (Fig 1B, down). The CV profile for all the EPSPs (Fig. 5D) for *in silico* (blue) is also closer to the *in vitro* (red) than the one computed applying UVR (Fig. 1D). Although our model has a slightly higher CV for the 6^th^, 7^th^ and 8^th^ EPSPs, the Kruskal-Wallis test showed no significant differences between both CV profiles for any of the EPSPs, demonstrating that the MVR hypothesis improved the synaptic variability of the circuit.

**Figure 5.**
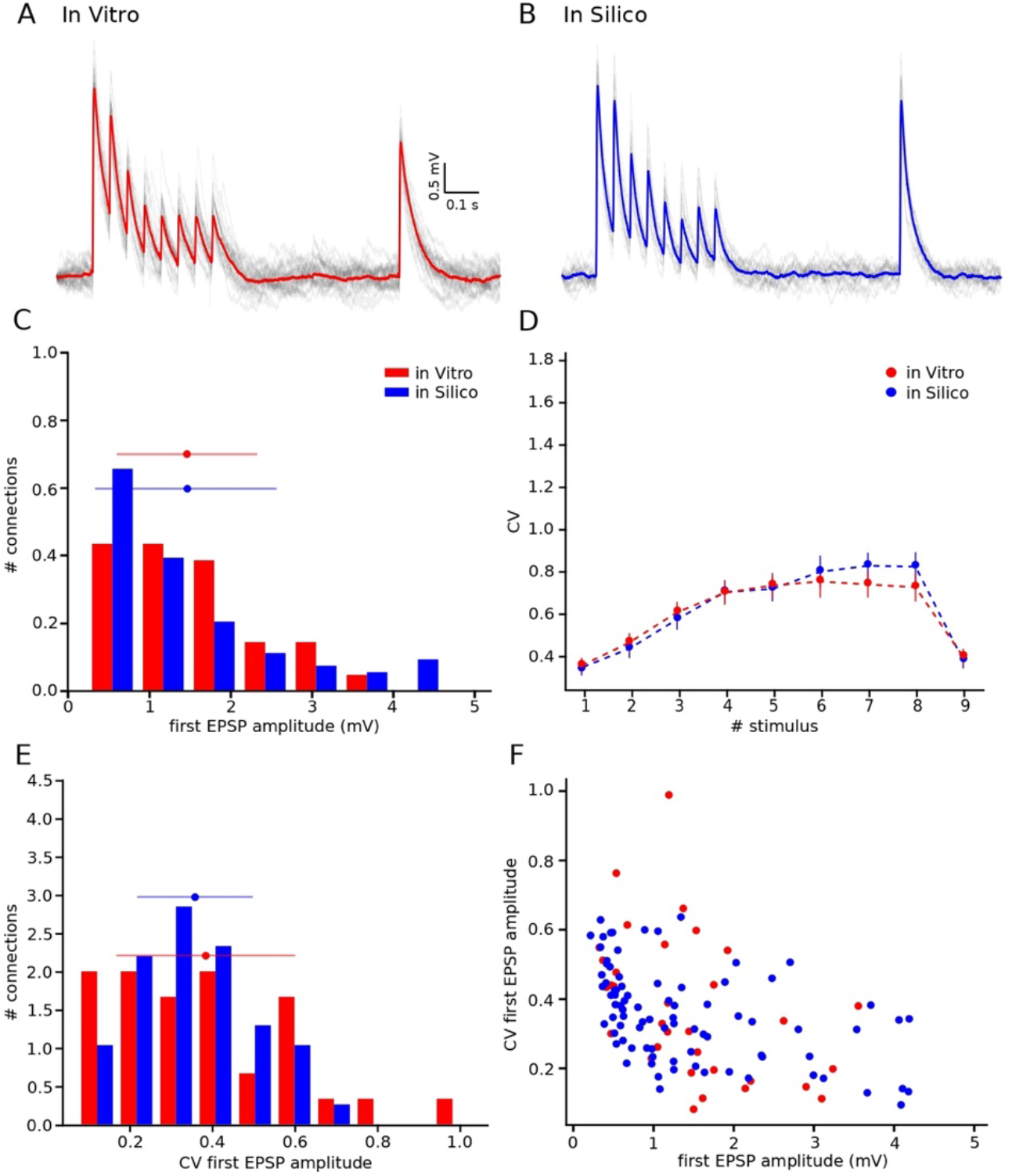
Releasing multiple vesicles improved the variability of the model. **(A)** *In vitro* mean voltage trace (red) of 20 protocol repetitions (grey) (same as in figure 1A down). **(B)** *In silico* mean voltage trace (blue) of 20 protocol repetitions (grey). **(C)** Distribution of the first EPSP amplitude for *in vitro* (red) and for the *in silico* (blue) experiments. **(D)** Mean CV profiles for the *in vitro* (red) and the *in silico* (blue) experiments. **(E)** CV Distribution of the first EPSP amplitude for *in vitro* (red) and the *in silico* (blue) data sets. **(F)** Raster plot of the first EPSP amplitude against the CV of the first EPSP amplitude for *in vitro* (red) and *in silico* (blue) experiments. All the *in silico* experiments are done with the N_RRP_ value that produces the minimum error. In the distributions and the CV profile, dots represent the mean and vertical and horizontal bars represent the standard deviation of all the experiments.

Another result that supported the improvement, was shown on the distributions for the first EPSP amplitude (Fig. 5C) and for the CV of the first EPSP amplitude (Fig. 5E). Both had a mean value closer than the one from the UVR like model as the statistic test showed no significant difference (mean EPSP values: 1.46 ± 0.86 mV for *in vitro*; 1.46 ± 0.95 mV for *in silico*; p = 0.69) (mean CV values: 0.38 ± 0.21 for *in vitro*; 0.35 ± 0.13 for *in silico*; p = 0.86). The distributions (Fig. 5C and E) were normalized to the respecting sample size such that the sum of products of width and height of each column is equal to the total count (33 for *in vitro*, 85 for *in silico*). Furthermore, the cross validated Kolmogorov-Smirnov test showed no significant difference between the first EPSP amplitude against the CV of the first EPSP amplitude for each data set (p = 0.29) (Fig. 5F), meaning that both data sets could come from the same population.

### 3.5 N_RRP_ prediction for other cell connections

Moreover, we extended this method to other cell connections for which we have models in our circuit, but no raw *in vitro* data sets. In this occasion we collected CVs data from the literature all of them found only for the 1^st^ EPSP amplitude.

Before computing the CV for the new cell connections, we had to consider that these CV were not computed using the JKK bootstrapping approach. From our previous analysis we know that the CV computed through the JKK method has a slightly different value than the CV computed without this mathematical approach. In the case of L5_TTPC connections *in vitro* data set the CV = 0.31 ± 0.14 while the CV_JKK_ = 0.38 ± 0.21 (CVs computed only for the first pulse), but the N_RRPs_ computed after 50 iterations in both cases were mostly the same with no significant difference between them (no-JKK N_RRP_ = 2.41 ± 1.08 and JKK N_RRP_ = 2.73 ± 1.22; p < 0.01) (Fig. 6A and B respectively). This value is smaller than the previous N_RRP_ obtained from comparing the CV for all the pulses, but it was expected as in the previous analysis we didn’t reach the exact CV value for the 1^st^ pulse, although there was no significant difference.

**Figure 6.**
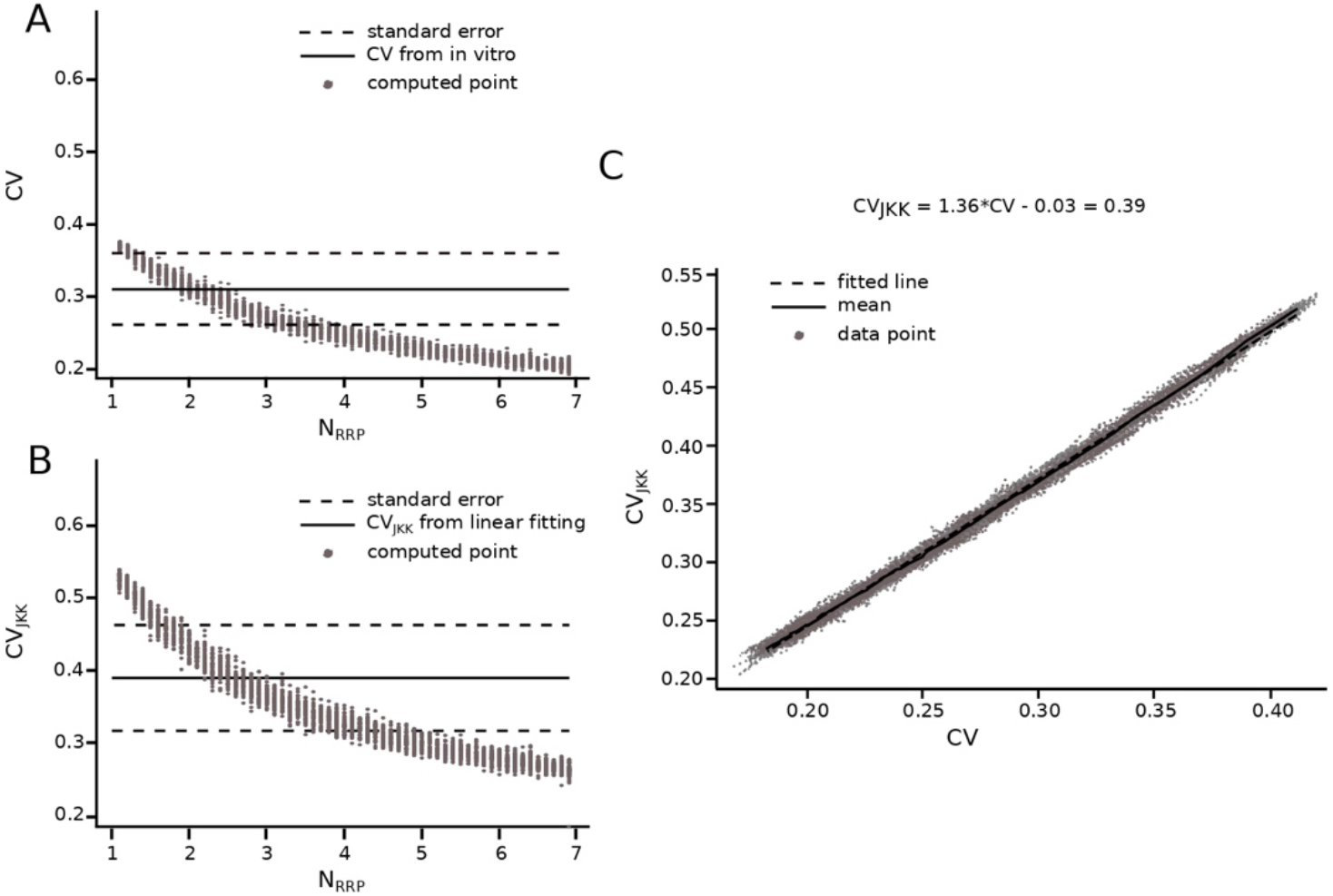
Extension of the method for connections reported in literature. Transformation from CV to CV_JKK_ using L5_TTPC connection as example **(A)** CV computed for different N_RRP_. Solid black line represents the CV computed for the *in vitro* data. Dotted black lines represent the standard error of the CV. **(B)** CV_JKK_ computed for different N_RRP_. Solid black line represents the CV_JKK_ obtained from the lineal fitting on **C**. Dotted black lines represent the standard error for this CV_JKK_. **(C)** CV to CV_JKK_ transformation. Solid black line represents the mean of the 50 iterations and dotted black line represent the linear fitting which equation is at the top of the plot. In **A** and **B** the grey dots show the 50 iterations from which we extract the best N_RRP_ as the one corresponding with the closest CV.

As we knew that the JKK bootstrapping method provide us with a more accurate manner for computing the EPSP amplitudes, we applied a transformation from CV to CV_JKK_ (Fig. 6C). We illustrate this transformation in figure 6 with L5_TTPC. First, we computed the CV of the first EPSP amplitude without (Fig. 6A) and with the JKK (Fig. 6B) analysis. Second, we represented both CVs in the same plot for the different N_RRP_ values and we fitted a line to the mean of the 50 repetitions (Fig. 6C). Once we obtained the linear equation we were able to determine the correspondent CV value computed with the JKK approach (Fig. 6B) for this connection we obtained CV_JKK_ = 0.39 ± 0.15 with a correspondent N_RRP_ = 2.84 ± 1.34. We did that for every connection for which we could find data in the literature and our simulation matched the variability (Table 1).

**Table 1.** Results for connections reported in literature. Table summarizing the CV_JKK_ computed for other five cell connections through the collection of data from literature and applying the JKK conversion explained in figure 6. For the L5_TTPC connection we used the CV computed in this work from our *in vitro* data set. L23_NBC_LBC: layer 2 and 3 nest and large basket cells; L23_PC: pyramidal cells in layer 2 and 3; L4_SSC: layer 4 spiny stellate cells; L5_TPC:C: thick tuft pyramidal cells that receive projections from thalamus; L5_SBC: small basket cells from layer 5.

| Connection type | Literature data | Jack-Knife conversion | Prediction |
| --- | --- | --- | --- |
| | CV | CV | $N_{RRP}$ |
| L23_NBC_LBC-L23_PC | $0.40 \pm 0.09$<br>(Wang, 2002) | $0.38 \pm 0.21$ | $1.96 \pm 0.98$ |
| L23_PC-L23_PC | $0.33 \pm 0.18$<br>(Feldmeyer, 2006) | $0.48 \pm 0.23$ | $2.60 \pm 1.28$ |
| L4_SSC-L23_PC | $0.27 \pm 0.13$<br>(Feldmeyer, 2002) | $0.37 \pm 0.09$ | $1.81 \pm 0.37$ |
| L4_SSC-L5_TPC:C | $0.33 \pm 0.20$<br>(Feldmeyer, 2005) | $0.46 \pm 0.15$ | $1.26 \pm 0.50$ |
| L5_TTPC-L5_SBC | $0.32 \pm 0.08$<br>(Wang, 2002) | $0.34 \pm 0.16$ | $1.82 \pm 0.90$ |
| L5_TTPC-L5_TTPC | $0.31 \pm 0.14$<br>(Our value) | $0.39 \pm 0.15$ | $2.84 \pm 1.34$ |

The results are summarized in Table 1 where we can observe that we were able to generalize for other five neuronal connections of the rat neocortex. We found that for layer 4 spiny stellate (L4_SSC) to layer 5 pyramidal cell connections that receive thalamic projections (L5_TPC:C) the transmission is mostly UV (N_RRP_ = 1.26 ± 0.50), while for the rest of the connections in the table the average number of vesicles ready to be released at the presynaptic terminal is between 2 to 3 (N_RRP_ = 2.60 ± 1.28 for L23_PC-L23_PC; N_RRP_ = 1.96 ± 0.98 for L23_NBC_LBC-L23_PC; N_RRP_ = 1.81 ± 0.37 for L4_SSC-L23_PC and N_RRP_ = 1.82 ± 0.90 for L5_TTPC-L5_SBC).

These results confirmed that implementing MVR in the model improved the synaptic dynamics with regard to the biological data. We found that for L5_TTPC connections N_RRP_ has variable value between 3 and 4 per synaptic contact. Furthermore, we predicted that MVR is most likely to happen in other cortical cell connections, supporting the idea that multiple vesicles are released in the neocortex influencing synaptic variability and information transmission.

## 4 Discussion

In this work we computed the N_RRP_ based on the previous work of Loebel and colleagues of 2009 (Loebel et al., 2009a) but we extended it to every active connection in a synapse. This approach is based on the comparison of the EPSPs amplitudes CV between a set of *in vitro* and *in silico* experiments with different N_RRP_ values performed in a data driven model of the rat somatosensory cortex (Markram et al., 2015) for a specific neuronal connection. The CV of the amplitude distributions is a currently used marker of the concentration of neurotransmitter in the synaptic cleft and for the postsynaptic receptor occupancy (Auger and Marty, 2000; Faber and Korn, 1991). For example, if a big quantity of neurotransmitter is released into the synaptic cleft then a high amplitude EPSP would be generated into the postsynaptic terminal. However, a large fraction of receptors would be occupied as well and consequently it would be more difficult to generate a second EPSP if more neurotransmitter is released. Thus, it is possible to measure the variability of the amplitude considering that high variability represents a small number of released vesicles.

### 4.1 UVR could not reproduce the variability observed into the biological data

Through this analysis we found that the UVR hypothesis did not reproduce the variability observed on the *in vitro* traces, in fact the CV profile for the *in silico* experiments was significantly larger, although the first EPSP amplitude was not statistically different. This result suggested us that the MVR hypothesis could be the appropriate approach to be considered for our model. On one hand, this idea differed from previous studies (Gulyás et al., 1993; Murphy et al., 2004; Redman, 1990) in which they claim that from each active contact in a synapse only one vesicle as most could be released, suggesting that the biological variability may come from changes in the quantal size. On the other hand, there were newer research studies that supported our idea of implementing the MVR hypothesis to improve our model (Brémaud et al., 2007; Hardingham et al., 2010; Huang et al., 2010; Loebel et al., 2009a; Rudolph et al., 2015). This discrepancy between works could come because they were studying different brain regions, with different experimental protocols, different species or different cell connections.

Before comparing both data collections, we extracted some important parameters from the *in vitro* group of data. First, we computed the parameters related with the deterministic model for short-term synaptic depression. For this purpose, we had to select only the depressing connections which 1^st^ EPSP amplitude was within the range of the *in vitro* data set and apply the deconvolution approach in order to be able to fit the peaks of each trace in the most precise possible manner. The values obtained were similar to values found in previous researches (Tsodyks and Markram, 1997b; Wang et al., 2006). Second, we calibrated the synaptic noise which represented the trial-to-trial variability into the synapse. Many studies support the idea that background synaptic noise is not only noise, but an addition of various meaningful procedures (Azouz and Gray, 1999; Faisal et al., 2008). Synaptic noise is classically shown in the spontaneous miniature postsynaptic currents, which is thought to be the result of spontaneous vesicle released (Fatt and Katz, 1950). This noise can influence not only the synaptic variability but also the transmission of information (Jacobson et al., 2005). Thus, while some studies did not support our idea of the contribution of the number of vesicles in the synaptic noise (Mackenzie et al., 2000), many others (Faisal et al., 2008; Franks et al., 2003; Pulido and Marty, 2017) encouraged us to complement our model with noise calibration.

### 4.2 L5_TTPC synapses are driven by multiple vesicles

In this research increasing the N_RRP_ improved the variability of the model, yielding synapses that behaved more similarly to the ones found in the biological system. Consequently, for cell connections between L5_TTPCs studied with a stimulation train at 20 Hz the resultant N_RRP_ mean was 3.78 ± 1.65 within a range between 1 to 9 vesicles. Considering that the number of active contacts found in L5_TTPC connections is between 4 to 8 (Markram et al., 1997), our prediction was that the range of total number of release sites for a synapse would be between 4 to 72. This estimation is in the range of previous studies in which they found that the range of the docked vesicles in L5_TTPC synapses was 2 – 30 (Rollenhagen and Lübke, 2006). Our predicted range is also consistent with previous estimates (Loebel et al., 2009a)(7 - 170), although in this case our range is considerably smaller. This difference could be due to the different approaches for the simulation: with the Montecarlo approach in the case of Loebel or with the data-driven model of the microcircuit in our case. Our result is also in accordance with a recent study in which they found that in L5B pyramidal cells of the rat somatosensory cortex at individual synaptic contacts the number of vesicles ready to be released ranged between 1.2 to 12.8 (Rollenhagen et al., 2018), although their mean value is slightly larger (5.40 ± 1.24). This difference could be due to the fact that we are considering all the TTPC in Layer 5 and not only the neurons in Layer 5B. Evidence from the mouse neuromuscular junction indicates a huge number of vesicles per active zone (~1700) (Ruiz et al., 2011). This large difference could be due to the stimulation frequency, which was 100 Hz comparing with our stimulation at 20 Hz, but it could be also due to the functionality. Synapses in the cortex probably need more variability as these areas are known to process information, while synapses between motoneurons and muscular fibers should be more reliable. Probably, pointing out the importance of MVR not only for the transmission of the information, but also as an important factor for defining the synaptic functionality.

The CV computed from the *in silico* experiments was slightly different without signification for the 6^th^, 7^th^ and 8^th^ EPSPs. We also found 1 vesicle of difference between the comparison with the train or comparing only the first pulse. It is known that many mechanisms are involved in vesicular release as the membrane fusion, receptor saturation, vesicle recycling or the barrier effect of the glia (Rizo and Xu, 2015; Rudolph et al., 2015; Stevens, 2003). We should take in consideration all these complex mechanisms for the future and complete our cortical model with them.

### 4.3 MVR also occurs in other cell connections of the rat neocortex

We extended the method to connections involving other cell types in the neocortex for which we had models and data from literature. We succeeded on five connections from which we concluded that on average the number of vesicles ready to be released is between 2 and 3. Although some studies did not support our result (Silver et al., 2003), many others support the idea of MVR as a process that happens in the neocortex (Zhang and Peskin, 2015) and actually that not necessarily the number of vesicles does not necessarily have to be the same, but that it could change from one layer to another (Brémaud et al., 2007). Other studies agreed with our result, suggesting that we achieve this result because the data that we were comparing with come from the somatosensory cortex (Huang et al., 2010). Consequently, it might be possible that MVR happens in some cortical areas but not in others, demonstrating that vesicle release is relevant for signal processing.

We had to extrapolate some data, for L5_TPC-L5_SBC we used the data from the work of Wang and colleagues of 2002 in which they describe the anatomy and synaptic behavior of basket interneurons in layers 2 – 4 (Wang et al., 2002). We took that decision since we could not find any reference from literature studying the necessary aspects of the synapse between pyramidal and basket cells in layer 5. If in the future data becomes available, it would be possible that the N_RRP_ for this connection changes. Another problem was that we did not manage to generalize all the connections for which we have models because we did not find any data in the literature or because the experimental data from the literature provided CV distributions with some very weak connections. In our model, even though we have weak connections, the CV distribution has a lower mean because most of the connections are stronger than those found in previous researches. Some studies provide a functional view of networks formed by weak connections as a manner to keep synchronicity in the brain (Bruno and Sakmann, 2006; Ren et al., 2017), showing the importance of the weak connections. We should study in depth this issue and come up with a solution in order to unravel the N_RRP_ for the rest of the connections.

In this work we provide a method to computed the N_RRP_ per active contact in a synapse. This method is an extension of Loebel and colleague’s work from 2009. Through the comparison between a set of *in vitro* and *in silico* data of the EPSP amplitude CV, we were able to find the correspondent N_RRP_. The results obtained by this method helped us to understand that MVR is a process that very likely happens in some cortical areas of the rat brain and that can be an important mechanism for the correct functioning of the information transmission through the brain.

## 6 Author Contributions

NB-Z developed and performed the data analysis and the *in silico* experiments, wrote the manuscript and illustrated the figures. JR developed and performed some data analysis. GC developed and performed some data analysis and some *in silico* experiments. RP designed and performed the *in vitro* experiments. HM critical revision of the manuscript. SR contributed to the writing and critical revision of the article. EM contributed to data interpretation, writing and critical revision of the manuscript.

